# Extreme heterogeneity in sex chromosome differentiation and dosage compensation in livebearers

**DOI:** 10.1101/589788

**Authors:** Iulia Darolti, Alison E. Wright, Benjamin A. Sandkam, Jake Morris, Natasha I. Bloch, Marta Farré, Rebecca C. Fuller, Godfrey R. Bourne, Denis M. Larkin, Felix Breden, Judith E. Mank

**Affiliations:** Department of Genetics, Evolution and Environment, University College London, UK; Department of Animal and Plant Sciences, University of Sheffield, UK; Department of Zoology, University of British Columbia, Canada; Department of Biomedical Engineering, University of Los Andes, Colombia; School of Biosciences, University of Kent, UK; Department of Animal Biology, University of Illinois at Urbana-Campaign, USA; Department of Biology, University of Missouri St. Louis, USA; Comparative Biomedical Sciences Department, Royal Veterinary College, UK; Department of Biological Science, Simon Fraser University, Canada; Department of Organismal Biology, Uppsala University, Sweden

## Abstract

Once recombination is halted between the X and Y chromosome, sex chromosomes begin to differentiate and transition to heteromorphism. While there is a remarkable variation across clades in the degree of sex chromosome divergence, far less is known about variation in sex chromosome differentiation within clades. Here, we combined whole genome and transcriptome sequencing data to characterise the structure and conservation of sex chromosome systems across Poeciliidae, the livebearing clade that includes guppies. We found that the *Poecilia reticulata* XY system is much older than previously thought, being shared not only with its sister species, *Poecilia wingei*, but also with *Poecilia picta*, which diverged 30 mya. Despite the shared ancestry, we uncovered an extreme heterogeneity across these species in the proportion of the sex chromosome with suppressed recombination, and the degree of Y chromosome decay. The sex chromosomes in *P. reticulata* are largely homomorphic, with recombination persisting over a substantial fraction. However, the sex chromosomes in *P. picta* are completely non-recombining and strikingly heteromorphic. ln addition to being highly divergent, the sex chromosome system in *P. picta* includes a neo-sex chromosome, the result of a fusion between the ancestral sex chromosome and part of chromosome 7. Remarkably, the profound degradation of the ancestral Y chromosome in *P. picta* is counterbalanced by the evolution of complete dosage compensation in this species, the first such documented case in teleost fish. Our results offer important insight into the initial stages of sex chromosome evolution and dosage compensation.

## INTRODUCTION

Sex chromosome evolution is characterized by remarkable variation across lineages in the degree of divergence between the X and Y chromosomes (1, 2). Derived from a pair of homologous autosomes, sex chromosomes begin to differentiate as recombination between them is supressed in the heterogametic sex over the region spanning a newly acquired sex-determining locus (3, 4). The lack of recombination exposes the sex-limited Y chromosome to a range of degenerative processes that cause it to diverge in structure and function from the corresponding X chromosome, which still recombines in females (5, 6). Consequently, the sex chromosomes are expected to eventually transition from a homomorphic to a heteromorphic structure, supported by evidence from many of the old and highly differentiated systems found in mammals (7, 8), birds (9), *Drosophila* (5) and snakes (10).

However, there is a significant heterogeneity among clades, and even among species with shared sex chromosome systems, in the spread of the non-recombining region, and the subsequent degree of sex chromosome divergence (11, 12). Age does not always reliably correlate with the extent of recombination suppression, as in some species the sex chromosomes maintain a largely homomorphic structure over long evolutionary periods (12-16), while in others the two sex chromosomes are relatively young, yet profoundly distinct (17). Comparing the structure and recombination patterns of sex chromosomes between closely related species is a powerful method to determine the forces shaping sex chromosome evolution over time.

Sex chromosome divergence can also lead to differences in X chromosome gene dose between males and females. Following recombination suppression, the Y chromosome undergoes gradual degradation of gene activity and content, leading to reduced gene dose in males (6, 18, 19). Genetic pathways that incorporate both autosomal and sex-linked genes are primarily affected by such imbalances in gene dose, with potential severe phenotypic consequences for the heterogametic sex (20). ln some species, this process has led to the evolution of chromosome-level mechanisms to compensate for the difference in gene dose (21, 22). However, the majority of sex chromosome systems are associated with gene-by-gene level mechanisms, whereby dosage sensitive genes are compensated, but overall expression of the X chromosome is lower in males compared to females (19, 22, 23).

As opposed to most mammals and birds, the sex chromosomes in many fish, lizard and amphibian species are characterized by a lack of heteromorphism, which has been attributed to processes such as sex chromosome turnover and sex reversal (15, 24-28). As a result, closely related species from these taxonomic groups often have a variety of sex chromosome systems found at different stages in evolution (26, 29). Additionally, global dosage compensation has not yet been found in fish, perhaps due to the transient nature of the sex chromosome systems and the general lack of heteromorphism in the group. However, incomplete dosage compensation, through a gene-by-gene regulation mechanism, may have evolved in sticklebacks (30) and flatfish (31).

Poeciliid species have been the focus of many studies concerning sex determination (25). Moreover, many poeciliids exhibit sexual dimorphism, with some colour patterns and fin shapes controlled by sex-linked loci (32-36). The clade also has a diversity of genetic sex determination systems, with both male and female heterogametic sex chromosomes observed in different species (37, 38). Most work on Poeciliid sex chromosome structure has focused on the *Poecilia reticulata* XY system, positioned on chromosome 12 (39) which shows very low levels of divergence (35, 40). Although recombination is suppressed over almost half the length of the *P. reticulata* sex chromosome, there is little sequence differentiation between the X and Y chromosomes and no perceptible loss of gene activity of Y-linked genes in males (40). This low level of divergence suggests a recent origin of the sex chromosome system.

There is intra-specific variation in the extent of the non-recombining region within *P. reticulata*, correlated with the strength of sexual conflict (40). Additionally, although *P. reticulata* and its sister species, *P. wingei*, are thought to share an ancestral sex chromosome system (41, 42), there is some evidence for variation in Y chromosome divergence between these species (42). lt is unclear whether the XY chromosomes maintain the same level of heteromorphism in other poeciliids (37, 41), or even whether they are homologous, to the sex chromosomes in *P. reticulata*.

Here we perform comparative genome and transcriptome analyses on multiple Poeciliid species to test for conservation and turnover of sex chromosome systems and investigate patterns of sex chromosome differentiation in the clade. We find the XY system in *P. reticulata* to be older than previously thought, being shared with both *P. wingei* and *P. picta* and thus dating back to at least 30 mya. Despite the shared ancestry, we uncover an extreme heterogeneity across these species in the size of the non-recombining region, the sex chromosomes being largely homomorphic in *P. reticulata* while completely non-recombining and highly diverged across the entire chromosome in *P. picta*. Additionally, we identify a fusion between the ancestral sex chromosome and part of chromosome 7 in *P. picta*, giving rise to a neo-sex chromosome. Remarkably, although the Y chromosome in *P. picta* shows signatures of profound sequence degeneration, we observe equal expression of X-linked genes in males and females, which we find to be the result of dosage compensation acting in this species. This is the first instance of complete sex chromosome dosage compensation reported in a fish.

## RESULTS AND DISCUSSION

### Comparative assembly of Poeciliid sex chromosomes

We sequenced the genome and transcriptome of three male and three female individuals from each of the four target species (*P. wingei, P. picta, P. latipinna* and *Gambusia holbrooki*) (Table S1), chosen to represent an even taxonomic distribution across Poeciliidae. For each species, we generated DNA-seq with an average of 222 million 150bp paired-end reads (average insert size 500bp, resulting in an average of 76X coverage), and 77.8 million 150bp mate-paired reads (average insert size 2kb, averaging 22X coverage) per individual. We also generated on average 26.6 million 75bp paired-end RNA-seq reads for each individual (see Materials and Methods). We combined this with published data from *P. reticulata* for comparative analyses (40).

Previous work on the sex chromosomes of these species showed evidence for male heterogametic systems in *P. reticulata* (40), *P. wingei* (41), *P. picta* (43) and *G. holbrooki* (44), and a female heterogametic system in *P. latipinna* (45, 46). For each target species, we built scaffold-level *de novo* genome assemblies using SOAPdenovo (47) (Table S2). Each assembly was constructed only using the reads from the homogametic sex in order to prevent co-assembly of X and Y reads. This allowed us to later assess patterns of sex chromosome divergence based on differences between the sexes in read mapping efficiency to the genome (detailed below).

To obtain scaffold positional information for each species, we used the Reference-Assisted Chromosome Assembly (RACA) algorithm (48), which integrates comparative genomic data, through pairwise alignments between the genomes of a target, an outgroup and a reference species, together with read mapping information from both sexes, to order target scaffolds into predicted chromosome fragments (see Materials and Methods, Table S2). RACA does not rely solely on sequence homology from the reference genome of *P. reticulata* (49) sex chromosome as a proxy for reconstructing the sex chromosomes in the target species, and instead incorporates read mapping and outgroup information as well from *Oryzias latipes* (50), which does not share the same sex chromosome pair. This approach is particularly important given the high rate of sex chromosome turnover observed in teleost fish (51), which could lead to substantial chromosomal rearrangement or other structural variation on sex chromosomes even across closely related species. lt also minimizes mapping biases that might result from different degrees of phylogenetic similarity of our target species to *P. reticulata*. Using RACA, we reconstructed chromosomal fragments in each target genome and identified syntenic blocks (regions which maintain sequence similarity and order) across the sex chromosomes of the target and the reference species. This provided both comparison at sequence level for each target species with reference genome and positional information of scaffolds in chromosome fragments.

### Extreme heterogeneity in sex chromosome differentiation patterns

In order to identify the extent of sex chromosome divergence in each of our target species, we first compared male and female read mapping rates (coverage) to the *de novo* scaffolds with chromosomal positional annotation from RACA. Regions that show substantial sequence differentiation between the two sex chromosomes result in reduced coverage in the heterogametic sex, as degeneration of the sex-limited chromosome leads to decreased mapping efficiency in that sex. We combined this with single nucleotide polymorphism (SNP) densities in males and females, which we expect to differ depending on the degree of similarity between the two sex chromosomes. ln sex-linked regions that have stopped recombining recently, but which still retain high sequence similarity to each other, we expect elevated SNP density in the heterogametic sex. ln male heterogametic systems, this is because Y reads still map to the female genome assembly, but carry many Y-specific mutations. ln regions where the Y has largely deteriorated, we expect SNP density to be lower in males compared to females, as highly diverged Y reads will not map to the female assembly, leaving the single hemizygous X haplotype. ln contrast, autosomal and pseudoautosomal regions are expected to show a similar coverage and SNP density between males and females.

Previous studies have suggested a very recent origin of the *P. reticulata* sex chromosome system on chromosome 12 based on its large degree of homomorphism and the limited expansion of the Y-specific region (40, 41). Contrary to these expectations, our combined coverage and SNP density analysis revealed that *P. reticulata, P. wingei* and *P. picta* share the same sex chromosome system (Fig. 1; Fig. S3; Fig. S4), revealing a shared ancestral system that dates back to at least 30 mya (52). Our findings suggest a far higher degree of sex chromosome conservation in this genus than we expected based on the small non-recombining region in *P. reticulata* in particular (40), and high degree of sex chromosome turnover in fish in general (53, 54). By contrast, in the *Xiphophorous* and *Oryzias* genera sex chromosomes have evolved independently between sister species (25, 55), and there are even multiple sex chromosomes within *X. maculatus* (56).

**Figure 1:**
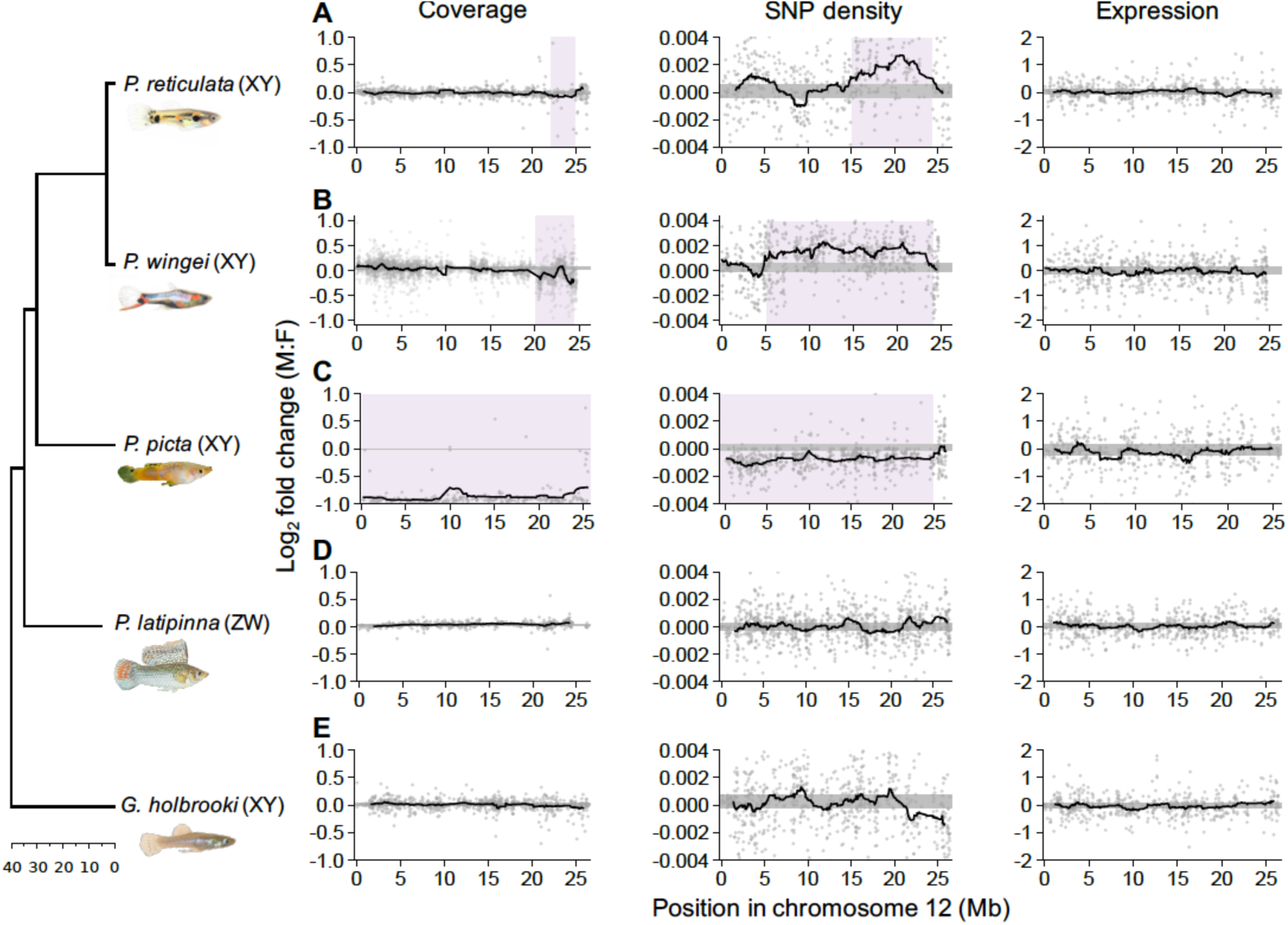
Differences between the sexes in coverage, SNP density and expression for *P. reticulata* chromosome 12 across all species. Moving average plots show male:female differences in sliding windows across the length of chromosome 12 in (A) *P. reticulata*, (B) *P. wingei*, (C) *P. picta*, (D) *P. latipinna* and (E) *G. holbrooki*. The 95% autosomal confidence intervals are shown in grey and regions of significant deviations from the 95% confidence intervals are highlighted in purple.

In addition to the unexpected conservation of this Poeciliid sex chromosome system, we observe extreme heterogeneity in patterns of X-Y recombination and differentiation across the three species. The *P. wingei* sex chromosomes have a similar, yet more accentuated, pattern of divergence compared to *P. reticulata* (Fig. 1A, B). The non-recombining region covers most of the *P. wingei* sex chromosomes and we can distinguish two evolutionary strata; an older stratum (20-25 Mb), showing significantly reduced male coverage, and a younger non-recombining stratum (5-20 Mb), as indicated by elevated male SNP density without a decrease in coverage (Fig. 1B). The old stratum has likely evolved ancestrally to *P. wingei* and *P. reticulata*, as its size and estimated level of divergence appear to be conserved in the two species. The younger stratum, however, has substantially expanded in *P. wingei* (40). These findings are consistent with the expansion of the heterochromatic block (41) and the large-scale accumulation of repetitive elements on the *P. wingei* Y chromosome (42).

More surprisingly, however, is the pattern of sex chromosome divergence that we recover in *P. picta*, which shows an almost 2-fold reduction in male to female coverage across the entire length of the ancestral sex chromosomes (Fig. 1C). This indicates not only that the ancestral Y chromosome in this species is completely non-recombining with the X, but also that the Y chromosome has undergone significant degeneration. Consistent with the notion that genetic decay on the Y will produce regions that are effectively hemizygous, we also recover a significant reduction in male SNP density (Fig. 1C).

In order to identify the ancestral Y region, we used *k*-mer analysis across *P. reticulata, P. wingei* and *P. picta* which detects shared male-specific *k*-mers, often referred to as Y-mers. Using this method, we have previously identified many shared male-specific sequences between *P. reticulata* and *P. wingei* (42) (Fig. S1A). Curiously, here we recovered very few shared Y-mers across all three species (Fig. S1A), which suggests two possible scenarios in the evolution of *P. picta* sex chromosomes. lt is possible that sex chromosome divergence began independently in *P. picta* compared to *P. reticulata* and *P. wingei*. Alternatively, the ancestral Y chromosome in *P. picta* may have been largely lost via deletion, resulting in either a very small Y chromosome, or an X0 system. To test for these alternative hypotheses, we reran the *k*-mer analysis in *P. picta* alone. We recovered almost twice as many female-specific *k*-mers than Y-mers in *P. picta* (Fig. S1A, B), which indicates that much of the Y chromosome is indeed missing. This is consistent with the coverage analysis (Fig.1C), which shows that male coverage of the X is half that of females, consistent with large-scale loss of homologous Y sequence.

From pairwise alignments we recovered a homologous syntenic block across all three species that spans the previously identified oldest evolutionary stratum of divergence on the *P. reticulata* sex chromosomes (22-25 Mb, Fig. S2) (40). We also identify a region of the X chromosome that is inverted in *P. reticulata* compared to in both *P. wingei* and *P. picta* (Fig. S2).

We also used differences in coverage and SNP density between males and females to identify the sex chromosomes in *P. latipinna* and *G. holbrooki*. We cannot identify any areas of divergence or restricted recombination on chromosome 12 of either of these species (Fig. 1D, E), indicating that the sex chromosomes in *P. reticulata, P. wingei* and *P. picta* formed after their common ancestor diverged from *P. latipinna*. Lack of conservation of the sex chromosome system on chromosome 12 is not unexpected for *P. latipinna*, as this species has evolved a female heterogametic system, however it implies that a different chromosome pair was recruited as the XY sex chromosome system in *G. holbrooki*.

We further attempted to find potential sex chromosome candidates in *P. latipinna* and *G. holbrooki* (Fig. S5, S6). While we can predict, based on coverage and SNP density, which chromosome pair is the likely sex chromosome candidate in each species (Fig. S5, S6), recombination across the genome shows significant local variation between the sexes and without some prior physical or genetic maps it is not possible to definitively determine the location of the sex-determining region. That said, the sex chromosomes in *P. latipinna* and *G. holbrooki* are either very young and undifferentiated, as would be expected given the high sex chromosome turnover in this clade, or have maintained an extremely homomorphic structure over relatively long evolutionary time.

### A neo-sex chromosome system in *Poecilia picta*

Interestingly, in *P. picta* we found strong support for a fusion between the ancestral sex chromosome (chromosome 12) and part of chromosome 7 (Fig. 2, Fig. S7; RACA adjacency = 0.52), giving rise to a neo-sex chromosome in this species. ln order to verify the fusion between chromosomes 12 and 7 in this species, we assessed the presence of bridge reads, with one read in each chromosomal segment, in our mate-pair data. We recovered a significantly higher number of mate-pair bridge reads between chromosome 12 and 7 than between chromosome 12 and any other chromosome (Fig. S7, Kruskal-Wallis, *p* < 0.001; pairwise Wilcoxon rank sum, all *p* < 0.007). We additionally excluded the possibility that our findings reflect a duplication of chromosome 7 into chromosome 12 instead of a fusion as the average coverage for this chromosome 7 region is similar to the average genomic coverage in both males (*p* = 0.09, Wilcoxon rank sum test) and females (*p* = 0.45, Wilcoxon rank sum test).

**Figure 2:**
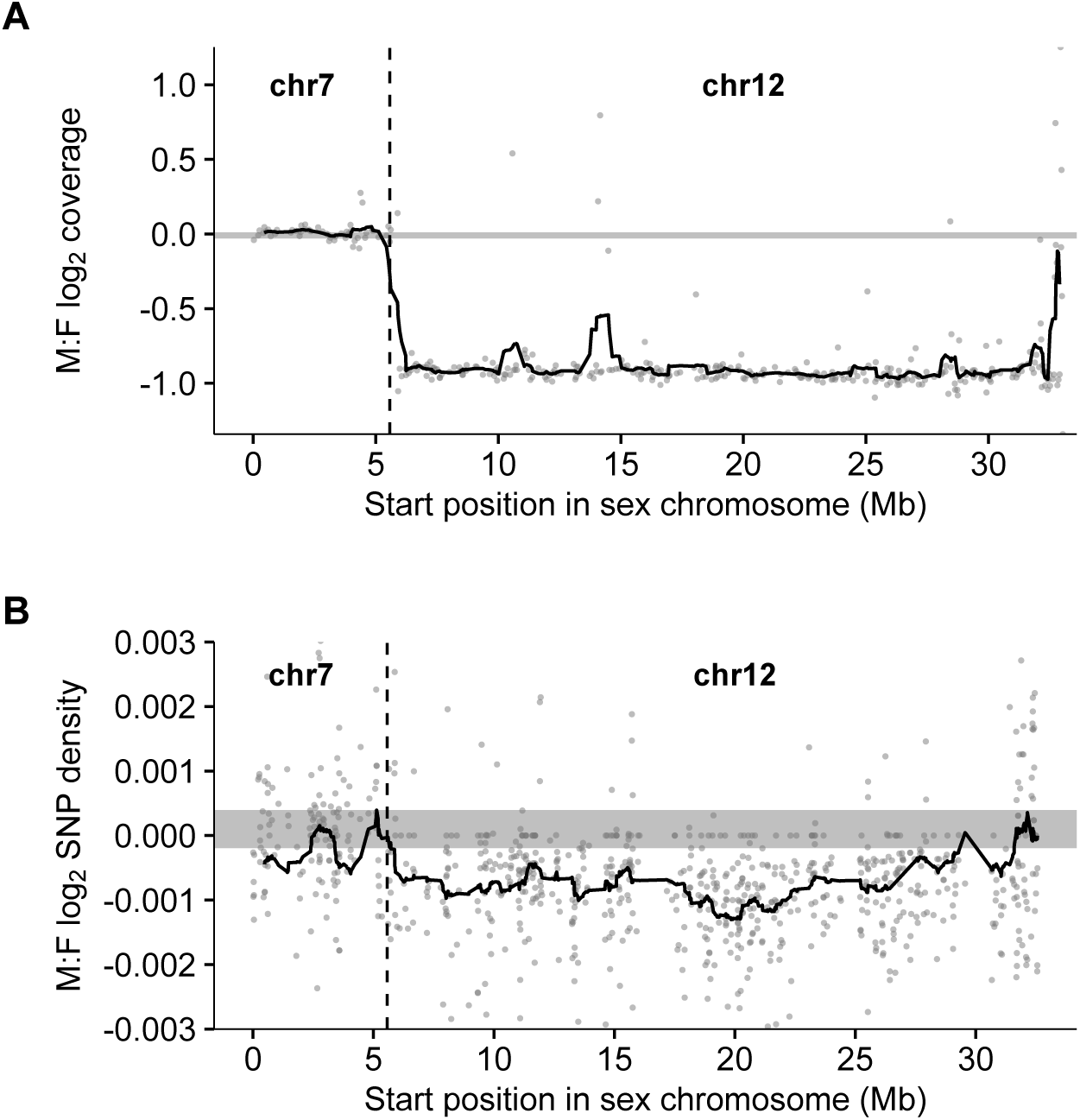
Coverage and SNP density characteristics of the *P. picta* sex chromosome system. Moving average of (A) coverage and (B) SNP density differences between the sexes in sliding windows across the neo-sex chromosome in *P. picta*. Shown in grey are the 95% confidence intervals based on bootstrapping autosomal values. The dotted vertical lines indicate the delimitation between the ancestral sex chromosome region (chr12) and the fused autosomal region (chr7).

The number of bridge reads supporting the fusion is similar in males (N=43) and females (N=39), suggesting that a very small region of the ancestral Y chromosome still remains and maintains some homology to the X near the fusion boundary. While the ancestral *P. picta* sex chromosomes are profoundly diverged, the fused autosomal region shows no differentiation in coverage or SNP density between the sexes (Fig. 2). This region of chromosome 7 thus comprises the only apparent pseudoautosomal region in this system.

In fishes, neo-sex chromosomes have been shown to facilitate phenotypic differentiation and speciation through reproductive isolation (26, 51, 57). Additionally, bringing specific sexually antagonistic loci in linkage with the sex-determining region may favour the fusion between an autosome and an existing sex chromosome (58, 59). Moreover, although *P. picta* inhabit a range of areas, from freshwater to brackish waters, there are regions where their distribution overlaps with that of *P. reticulata* (60). Both species exhibit similar life-history traits (60), however there is evidence of reproductive barriers between them (61). On the other hand, hybridization between *P. reticulata* and *P. wingei*, for example, occurs frequently in areas of sympatry (62). lt is possible that the neo-sex chromosome system in *P. picta* facilitates reproductive isolation, however, further studies would be needed to test this.

### Y degeneration and dosage compensation in *Poecilia picta*

In order to investigate the extent of Y gene activity decay in our target species, we assessed differences in gene expression between the sexes, and between the autosomes and the non-recombining regions of the sex chromosomes (Fig. 3D). Consistent with the high degree of sequence conservation between the X and the Y chromosomes in *P. reticulata* and *P. wingei*, we find no decrease in male gene expression in these species. Surprisingly, while our results indicate Y chromosome degeneration in *P. picta*, here also we did not find a significant reduction in average male:female gene expression on the sex chromosomes compared to the autosomes in males (*p* = 0.92, Wilcoxon rank sum test), despite the slightly lower gene expression on the sex chromosomes in males compared to females (*p* = 0.002, Wilcoxon signed rank test) (Fig. 3D). This finding suggests the action of a chromosome-level compensation mechanism in *P. picta* to counteract the imbalance in gene dose in males.

**Figure 3:**
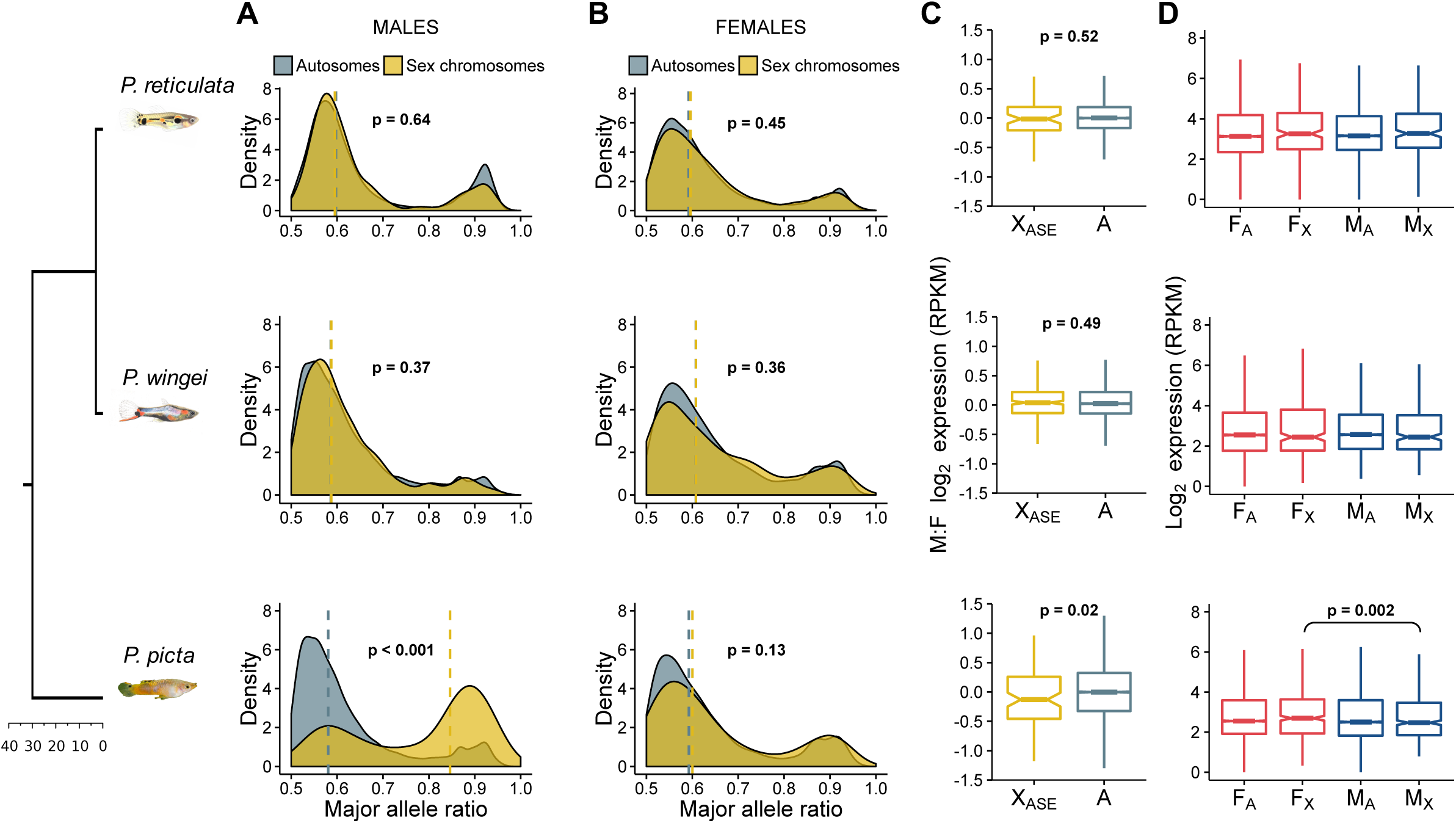
Patterns of gene and allele-specific expression. Density plots show the distribution of the major allele frequency in autosomes (grey) and sex chromosomes (yellow) in (A) males and (B) females of each species. Vertical dotted lines indicate median values and *p* values are based on Wilcoxon rank sum tests. (C) Boxplots show differences in expression between the sexes for autosomal genes (grey) and sex chromosome genes with an ASE pattern in males (yellow). *P* values are based on Wilcoxon rank sum tests. (D) Boxplots show average male and female expression for autosomes and the non-recombining region of the sex chromosomes. The *p* value is based on Wilcoxon signed rank test.

To further investigate the presence of dosage compensation, we estimated allele-specific expression patterns (ASE) (Fig. 3). The diverged Y chromosome in *P. picta* should be reflected in a significantly unbalanced contribution from the two sex chromosomes in males to the overall expression of heterozygous sites. We found that, on average, only 128 (24%) of the genes on the X chromosome in *P. picta* have SNPs in males, while 239 (45%) have SNPs in females. The large proportion of hemizygous genes in males is consistent with the large-scale deletion of much of the Y chromosome in *P. picta*, which is also coherent with our *k*-mer analysis.

Of the 128 X-linked male-heterozygous genes, 110 show significant allele-specific expression (Fig. 3A). lndeed, our allelic differential expression analysis revealed that a significantly larger proportion of heterozygous sites show an ASE pattern on the sex chromosomes than on the autosomes in *P. picta* males (x^2^ = 70.565, *p* < 0.001, Chi-square test; Fig. 3A). ln contrast, in *P. picta* females and in both males and females of *P. reticulata* and *P. wingei*, the majority of heterozygous sites throughout the genome show equal transcription between the maternal and paternal chromosomes (Fig. 3A, B).

Remarkably, despite being expressed from a single active gene copy in males, these ASE genes had a broadly similar, yet significantly higher (*p* = 0.02, Wilcoxon rank sum test), difference in expression between males and females, compared to autosomal genes (Fig. 3C). The marginally lower expression of single-copy X-linked genes in males likely represents functional dosage compensation, and is far more effective than the partial, localised, dosage compensation recorded in other fish species (30, 31).

### Concluding remarks

Our comparative analyses reveal a striking heterogeneity in the degree of recombination suppression and Y chromosome degeneration across Poeciliid species with a shared sex chromosome system. Through multiple independent lines of evidence, including sequence coverage, *k*-mer analysis and ASE patterns, we show a profound degeneration of the Y chromosome in *P. picta*. Remarkably, the hemizygosity of the X in males has led to the evolution of complete dosage compensation in this species, the first such instance found across teleost fish. Additionally, the sex chromosome system in *P. picta* contains a neo-sex chromosome, from the fusion of the ancestral sex chromosome with an autosome. Our findings highlight the importance of comparative studies of sex chromosome variation within clades and suggest that fish may harbour extensive variation in sex chromosome evolution.

## MATERIALS AND METHODS

### Sample collection and sequencing

We collected adult male and female individuals from four guppy species (*Poecilia wingei* from our laboratory population, *Poecilia picta* from Guyana, *Poecilia latipinna* and *Gambusia holbrooki* from Florida, USA). We chose these samples in order to obtain an even phylogenetic distribution.

All samples were collected in accordance within ethical guidelines. *P. latipinna* and *G. holbrooki* were collected under Florida permit FNW17-10 and St. Mark’s Refuge permit FF04RFSM00-17-09. *P. picta* was collected under permit from the Environmental Protection Agency of Guyana (Permit 120616 SP: 015). *P. wingei* was collected from our lab population, a colony of a strain maintained by a UK fish fancier.

From each species, we immediately stored head and tail samples from three males and three females in ethanol and, RNAlater, respectively. We extracted DNA from heads with the DNeasy Blood and Tissue Kit (Qiagen) and RNA from tails with the RNeasy Kit (Qiagen), following the manufacturer’s instructions. Library preparation and sequencing were performed at The Wellcome Trust Centre for Human Genetics, University of Oxford, following standard protocols and using the lllumina HiSeq 4000 platform. Genomic DNA was used to construct both short-insert (average insert size 500bp) and long-insert (average insert size 2kb) sequencing libraries for each individual. We assessed data quality with FastQC v0.11.3 (www.bioinformatics.babraham.ac.uk/projects/fastqc/) and used Trimmomatic v0.36 (63) to trim reads. For both DNA-seq and RNA-seq reads we removed adaptor sequences, regions of low Phred score (reads with average Phred score <15 in sliding windows of four bases and reads with leading/trailing bases with a Phred score < 3) and short reads (if either read in a pair was shorter than 50bp).

### Genome assembly

We first corrected the reads using Quake v0.3.5 (64) and estimated the optimal assembly *k*-mer length using KmerGenie v1.6741 (65). We then used SOAPdenovo v2.04 (47) to construct female *de novo* genome assemblies for *P. wingei, P. picta* and *G. holbrooki* and a male assembly for *P. latipinna*, using both the paired-end and mate-pair reads (Table S2).

The paired-end reads were used for both the contig and scaffolding steps of the assembly process, while the mate-pair reads were only used for scaffolding. Additionally, we used GapCloser v1.12 to close the gaps resulting from the assembly scaffolding step. Finally, we removed sequences shorter than 1kb from the assemblies.

To improve assembly contiguity and reconstruct chromosomal fragments for each species, we followed the UCSC chains and nets pipeline from the kentUtils software suite (66) before employing the Reference-Assisted Chromosome Assembly (RACA) algorithm (48). The chains and nets pipeline is designed for building pairwise nucleotide alignments and bridging gaps between pairwise syntenic blocks to construct larger structures (66). A chain alignment represents an ordered pairwise sequence alignment between two species. A net alignment represents a collection of chains within a genome region, ordered in a hierarchical manner based on synteny scoring. The RACA algorithm incorporates the pairwise alignment files, together with read mappings information to identify syntenic fragments (regions which maintain sequence similarity and order) across the species used. RACA then estimates adjacency between syntenic fragments in each target genome to reconstruct predicted chromosome fragments (PCFs) for each target species (48).

Firstly, for each species, we carried out DNA-seq read mappings to the *de novo* assemblies using Bowtie2 v2.3.3.1 (67), reporting concordant mappings only (--no-discordant option) and using the appropriate mate orientations according to the insert size of the libraries (--fr option for short-insert libraries and -rf option for long-insert libraries). The resulting alignments were converted into the RACA-specific input format (script available on the RACA website http://bioen-compbio.bioen.illinois.edu/RACA/).

We also obtained pairwise alignments using LASTZ (www.bx.psu.edu/%7Ersharris/lastz/; parameters C=0, E=30, H=2000, K=3000, L=3000, O=400, M=50) between a reference species (here we used the *P. reticulata* genome, obtained from NCBl GenBank Guppy_female_1.0+MT, assembly accession GCF_000633615.2), the target species and an outgroup species (here we used the Medaka, *Oryzias latipes*, genome, obtained from GenBank ASM223467v1, assembly accession GCA_002234675.1). We then converted these alignments into chains and nets formats following the UCSC axtChain (-minScore=1000, - linearGap=medium), chainAntiRepeat, chainSort, chainPreNet and netSyntenic tools (66). The syntenic chains and nets fragments, together with the paired-end alignments, were used as input files for RACA (resolution=10000 for *P. picta* and *P. latipinna* and resolution=1000 for *P. wingei* and *G. holbrooki*). For each target species, RACA ordered and oriented target scaffolds into PCFs (Table S2), and we used this positional information of scaffolds in the genome for all further analyses.

### Analysis of genomic coverage

For each species, using BWA v0.7.12 (68), we mapped male and female paired-end DNA-seq reads to the *de novo* scaffolds with positional annotation from RACA, following the aln and sampe alignment steps, and extracted uniquely mapping reads. We then used soap.coverage v2.7.9 (http://soap.genomics.org.cn/) to calculate the coverage (number of times each site was sequenced divided by the total number of sequenced sites) of each scaffold in each sample. For each scaffold, we calculated fold change coverage (male:female) as log_2_(average male coverage) - log_2_(average female coverage).

### SNP density analysis

For each species, using Bowtie1 v1.1.2 (67), we mapped male and female paired-end DNA-seq reads to the *de novo* scaffolds with positional annotation from RACA, generating map format output files. We sorted the map files by scaffold and converted them into profiles, which represent counts for each of the four nucleotide bases, for each individual using bow2pro v0.1 (http://guanine.evolbio.mpg.de/). For each site, we applied a minimum coverage threshold of 10 and called SNPs as sites with a major allele frequency of 0.3x the total site coverage. We obtained gene information through the expression analysis detailed below and for each gene we calculated the average SNP density as the number of SNPs divided by the number of filtered sites. We excluded SNPs outside of genic regions. For each gene we then calculated fold change SNP density (male:female) as log_2_(average male SNP density) - log_2_(average female SNP density).

### Gene expression analysis

For each species, using HlSAT2 v2.0.4 (69), we mapped male and female RNA-seq reads to the *de novo* scaffolds with positional annotation from RACA, reporting paired (--no-mixed) and concordant (--no-discordant) mappings only and tailoring the alignments for downstream transcript assembly (--dta). We used SAMtools to coordinate sort and bam convert the sam output files. For each sample, we then used StringTie (70) to obtain transcripts in a GTF file format, which we then merged to assemble a non-redundant set of transcripts for each species. Before further analyses, we filtered the merged GTF file for non-coding RNA (ncRNA) by using BEDtools getfasta (71), extracted target transcript sequences and removed transcripts with BLAST hit to ncRNA sequences from *Poecilia formosa* (PoeFor_5.1.2), *Oryzias latipes* (MEDAKA1), *Gasterosteus aculeatus* (BROADS1), and *Danio rerio* (GRCz10), obtained from Ensembl 84 (72).

For each species, we estimated gene expression by extracting read counts for each gene using HTSeq-count (73) and the ncRNA filtered transcriptome. We only kept genes that were placed on scaffolds with positional information on PCFs. For these genes, we converted read counts to RPKM values with edgeR (74), normalised with TMM, and applied a minimum expression threshold of 2RPKM in half or more of the individuals in one sex. For each gene we then calculated fold change expression (male:female) as log_2_(average male expression) - log_2_(average female expression).

### *k*-mer analysis

In order to identify shared Y sequence across *P. reticulata, P. wingei* and *P. picta*, we followed a *k*-mer analysis method previously described in Morris et al. 2018 (42). We have previously used this approach to successfully identify shared Y sequence between *P. reticulata* and *P. wingei* (42). Briefly, here we used the HAWK pipeline (75) to count *k*-mers from paired-end DNA-seq reads and identify unique *k*-mers for each sex in each species. Across all the species, we then identified shared female unique *k*-mers and shared male unique *k*-mers, referred to as Y-mers (42).

### Allele-specific expression (ASE) analysis

In order to estimate ASE patterns from RNA-seq data, we tailored previously published pipelines (76, 77). For each species, we called SNPs separately for males and females using Samtools mpileup and varscan (78), with parameters --min-coverage 2, --min-ave-qual 20, -- min-freq-for-hom 0.90, and excluding triallelic SNPs and Ns. Additionally, we removed SNPs that were not located within genic regions from the final filtered gene dataset. To exclude potential sequencing errors from our SNP dataset, we applied coverage filtering thresholds (76, 77). Firstly, we set a minimum site coverage of 15 reads (the sum of major and minor alleles), as a power analysis indicated that at a minimum coverage of 15 reads we have a 78% power to detect a signal of allele-specific expression. Secondly, we applied a variable coverage filter that accounts for the change in the likelihood of sequencing errors at different coverage levels (accounting for an error rate of 1 in 100 and a maximum coverage for a given site of 100,000 (76)). Lastly, to avoid the potential bias in our ASE estimations from the preferential assignment of reads to the reference allele (79), we removed clusters of more than 5 SNPs in 100 bp windows.

If genes have biallelic expression, meaning that alleles from both chromosomes are expressed at the same level, we expect a probability of around 0.5 of recovering reads from either chromosome. For each SNP in the final filtered dataset we tested for ASE by identifying significant deviations from the expected probability of 0.5 using a two-tailed binomial test (*p* < 0.05). We corrected for multiple testing when running binomial tests on autosomal SNPs. Additionally, we called SNPs as ASE if a minimum of 70% of the reads stemmed from one of the chromosomes. We called genes as ASE if they had at least one SNP with a consistent ASE pattern across all heterozygous samples. We tested for significant differences in ASE patterns between the sexes and between the autosomes and the sex chromosomes using chi-square tests.

## ACKNOWLEDGEMENTS

This work was supported by the European Research Council (grant agreements 260233 and 680951 to J.E.M.), the Biotechnology and Biological Sciences Research Council (PhD to l.D., grant number BB/M009513/1). CElBA Biological Center partially subsidized our expenses during field collection in Guyana. We acknowledge the use of the UCL Legion High Performance Computing Facility (Legion@UCL) in the completion of this work. We thank P. Almeida, A. Corral-Lopez, B. Furman, D. Metzger, J. Shu and W. van der Bijl for helpful comments and suggestions.

## AUTHOR CONTRIBUTIONS

J.E.M. and l.D. designed the research; B.S., R.F., G.R.B., F.B. and J.E.M. collected the samples; G.R.B. provided logistical support in Guyana; l.D. performed DNA and RNA extractions; N.l.B. performed RNA extractions for *P. wingei*; l.D. and A.E.W. performed the research; J.M. performed the *k*-mer analysis; l.D. analysed the data; M.F. and D.M.L. provided support with the chains and nets pipeline and the RACA algorithm; l.D. and J.E.M. wrote the manuscript; all authors revised the manuscript.

## DATA ACCESSIBILITY

DNA-seq and RNA-seq reads have been deposited at the NCBl Sequencing Read Archive (BioProject lD PRJNA353986 for *P. reticulata* reads; PRJNA528814 for *P. wingei, P. picta, P. latipinna* and *G. holbrooki* reads) and at the European Nucleotide Archive (lD PRJEB26489 for *P. wingei* paired-end DNA-seq reads).

## REFERENCES

1. Bachtrog D, Kirkpatrick M, Mank JE, McDaniel SF, Pires JC, Rice W,Valenzuela N (2011) Are all sex chromosomes created equal? Trends Genet 27(9):350-357.

2. Bachtrog D, Mank JE, Peichel CL, Kirkpatrick M, Otto SP, Ashman TL, Hahn MW, Kitano J, Mayrose I, Ming R, Perrin N, Ross L, Valenzuela N, Vamosi JC, Tree of Sex C (2014) Sex determination: why so many ways of doing it? PLoS Biol 12(7):e1001899.

3. Muller HJ (1918) Genetic variability, twin hybrids and constant hybrids, in a case of balanced lethal factors. Genetics 3:422.

4. Ohno S (1967) Sex chromosomes and sex-linked genes (Springer, New York).

5. Bachtrog D (2013) Y-chromosome evolution: emerging insights into processes of Y- chromosome degeneration. Nat Rev Genet 14(2):113-124.

6. Charlesworth B,Charlesworth De (2000) The degenration of Y chromosomes. Philosophical Transactions of the Royal Society of London. Series B: Biological Sciences 355:1563–1572.

7. Lahn BT,Page DC (1999) Four evolutionary strata on the human X chromosome. Science 286:964–967.

8. Skaletsky H, Kuroda-Kawaguchi T, Minx PJ, Hillier HSC, Brown LG, Repping S, Pyntikova T, Ali J, Bieri T,Chinwalla A (2003) The male-specific region of the human Y chromosome is a mosaic of discrete sequence classes. Nature 423:825.

9. Wright AE, Harrison PW, Montgomery SH, Pointer MA,Mank JE (2014) Independent stratum formation on the avian sex chromosomes reveals interchromosomal gene conversion and predominance of purifying selection on the W chromosome. Evolution 68(11):3281–3295.

10. Matsubara K, Tarui H, Toriba M, Yamada K, Nishida-Umehara C, Agata K,Matsuda Y (2006) Evidence for different origin of sex chromosomes in snakes, birds, and mammals and step-wise differentiation of snake sex chromosomes. PNAS 28:18190– 18195.

11. Fujito S, Takahata S, Suzuki R, Hoshino Y, Ohmido N,Onodera Y (2015) Evidence for a Common Origin of Homomorphic and Heteromorphic Sex Chromosomes in Distinct Spinacia Species. G3 (Bethesda) 5(8):1663–1673.

12. Vicoso B, Emerson JJ, Zektser Y, Mahajan S,Bachtrog D (2013) Comparative sex chromosome genomics in snakes: differentiation, evolutionary strata, and lack of global dosage compensation. PLoS Biol 11(8):e1001643.

13. Xu L, Sin SYW, Grayson P, Janes DE, Edwards SV,Sackton TB (2018).

14. Vicoso B,Bachtrog D (2013) Sex-biased gene expression at homomorphic sex chromosomes in emus and its implication for sex chromosome evolution. PNAS 110:6453–6458.

15. Stock M, Horn A, Grossen C, Lindtke D, Sermier R, Betto-Colliard C, Dufresnes C, Bonjour E, Dumas Z, Luquet E,Maddalena T (2011) Ever-young sex chromosomes in European tree frogs. PLoS Biol 9:e1001062.

16. Ahmed S, Cock JM, Pessia E, Luthringer R, Cormier A, Robuchon M, Sterck L, Peters AF, Dittami SM, Corre E,Valero M (2014) A haploid system of sex determination in the brown alga Ectocarpus sp. Curr. Biol. 8:1945–1957.

17. Bergero R, Forrest A, Kamau E,Charlesworth D (2007) Evolutionary strata on the X chromosomes of the dioecious plant Silene latifolia: evidence from new sex-linked genes. Genetics 175:1945–1954.

18. Charlesworth B (1978) Model for evolution of Y chromosomes and dosage compensation. Proc Natl Acad Sci U S A 75(11):5618–5622.

19. Mank JE (2009) The W, X, Y and Z of sex-chromosome dosage compensation. Trends in Genetics 25(5):226–233.

20. Malone JH, Cho DY, Mattiuzzo NR, Artieri CG, Jiang L, Dale RK, Smith HE, McDaniel J, Munro S, Salit M, Andrews J, Przytycka TM,Oliver B (2012) Mediation of Drosophila autosomal dosage effects and compensation by network interactions. Genome Biol 13:R28.

21. Mank JE, Hosken DJ,Wedell N (2011) Some inconvenient truths about sex chromosome dosage compensation and the potential role of sexual conflict. Evolution 65(8):2133–2144.

22. Mullon C, Wright AE, Reuter M, Pomiankowski A,Mank JE (2015) Evolution of dosage compensation under sexual selection differs between X and Z chromosomes. Nat Commun 6:7720.

23. Mank JE (2013) Sex chromosome dosage compensation: definitely not for everyone. Trends Genet 29(12):677–683.

24. Ezaz T, Sarre SD, O’Meally D, Graves JA,Georges A (2009) Sex chromosome evolution in lizards: independent origins and rapid transitions. Cytogenet Genome Res 127(2-4):249-260.

25. Volff JN,Schartl M (2001) Variability of genetic sex determination in poeciliid fishes. Genetica 111(1-3):101–110.

26. Ross JA, Urton JR, Boland J, Shapiro MD,Peichel CL (2009) Turnover of sex chromosomes in the stickleback fishes (gasterosteidae). PLoS Genet 5(2):e1000391.

27. Dufresnes C, Borzee A, Horn A, Stock M, Ostini M, Sermier R, Wassef J, Livinchuck SN, Kosch TA, Waldman B,Jang Y (2015) Sex-chromosome homomorphy in Palearctic tree frogs results from both turnovers and X–Y recombination. Mol Biol Evol 8:2328–2337.

28. Stock M, Savary R, Betto-Colliard C, Biollay S, Jourdan-Pineau H,Perrin N (2013) Low rates of X-Y recombination, not turnovers, account for homomorphic sex chromosomes in several diploid species of Palearctic green toads (Bufo viridis subgroup). J Evol Biol 26(3):674–682.

29. Cnaani A, Lee BY, Zilberman N, Ozouf-Costaz C, Hulata G, Ron M, D’Hont A, Baroiller JF, D’Cotta H, Penman DJ,Tomasino E (2008) Genetics of sex determination in tilapiine species. Sexual Development 2:43–54.

30. Leder EH, Cano JM, Leinonen T, O’Hara RB, Nikinmaa M, Primmer CR,Merila J (2010) Female-biased expression on the X chromosome as a key step in sex chromosome evolution in threespine sticklebacks. Mol Biol Evol 27(7):1495–1503.

31. Chen S, Zhang G, Shao C, Huang Q, Liu G, Zhang P, Song W, An N, Chalopin D, Volff JN, Hong Y, Li Q, Sha Z, Zhou H, Xie M, Yu Q, Liu Y, Xiang H, Wang N, Wu K, Yang C, Zhou Q, Liao X, Yang L, Hu Q, Zhang J, Meng L, Jin L, Tian Y, Lian J, Yang J, Miao G, Liu S, Liang Z, Yan F, Li Y, Sun B, Zhang H, Zhang J, Zhu Y, Du M, Zhao Y, Schartl M, Tang Q,Wang J (2014) Whole-genome sequence of a flatfish provides insights into ZW sex chromosome evolution and adaptation to a benthic lifestyle. Nat Genet 46(3):253–260.

32. Lindholm A,Breden F (2002) Sex chromosomes and sexual selection in poeciliid fishes. Am Nat 160 Suppl 6:S214–224.

33. Gordon SP, Lopez-Sepulcre A,Reznick DN (2012) Predation-associated differences in sex linkage of wild guppy coloration. Evolution 66(3):912–918.

34. Winge Ö (1927) The location of eighteen genes in Lebistes reticulatus. Journal of Genetics 18:1–43.

35. Tripathi N, Hoffmann M, Willing EM, Lanz C, Weigel D,Dreyer C (2009) Genetic linkage map of the guppy, Poecilia reticulata, and quantitative trait loci analysis of male size and colour variation. Proc Biol Sci 276(1665):2195–2208.

36. Lindholm AK, Brooks R, Breden F (2004) Extreme polymorphism in a Y-linked sexually selected trait. Heredity (Edinb) 92(3):156–162.

37. Schultheis C, Bohne A, Schartl M, Volff JN,Galiana-Arnoux D (2009) Sex determination diversity and sex chromosome evolution in poeciliid fish. Sex Dev 3(2-3):68-77.

38. Traut W,Winking H (2001) Meiotic chromosomes and stages of sex chromosome evolution in fish: zebrafish, platyfish and guppy. Chromosome Res 9(8):659–672.

39. Tripathi N, Hoffmann M, Weigel D,Dreyer C (2009) Linkage analysis reveals the independent origin of Poeciliid sex chromosomes and a case of atypical sex inheritance in the guppy (Poecilia reticulata). Genetics 182(1):365–374.

40. Wright AE, Darolti I, Bloch NI, Oostra V, Sandkam B, Buechel SD, Kolm N, Breden F, Vicoso B,Mank JE (2017) Convergent recombination suppression suggests role of sexual selection in guppy sex chromosome formation. Nat Commun 8:14251.

41. Nanda I, Schories S, Tripathi N, Dreyer C, Haaf T, Schmid M,Schartl M (2014) Sex chromosome polymorphism in guppies. Chromosoma 123(4):373–383.

42. Morris J, Darolti I, Bloch NI, Wright AE, Mank JE (2018) Shared and Species-Specific Patterns of Nascent Y Chromosome Evolution in Two Guppy Species. Genes (Basel) 9(5).

43. Lindholm AK, Sandkam B, Pohl K,Breden F (2015) Poecilia picta, a Close Relative to the Guppy, Exhibits Red Male Coloration Polymorphism: A System for Phylogenetic Comparisons. PLoS One 10(11):e0142089.

44. Russo C, Rocco L, Stingo V, Aprea G,Odierna G (2009) A cytogenetic analysis ofGambusia holbrooki(Cyprinodontiformes, Poecilidae) from the River Sarno. Italian Journal of Zoology 66(3):291–296.

45. Sola JL, Rossi AR, Iaselli V, Rasch EM,Monaco PJ (1992) Cytogenetics of bisexual/unisexual species of Poecilia. Cytogenet Genome Res 60:229–235.

46. Sola L, Bressanello S, Rasch EM,Monaco PJ (1993) Cytogenetics of bisexual/unisexual species of Poecilia. IV. Sex chromosomes, sex chromatin composition and Ag-NOR polymorphisms in Poecilia latipinna: a population from Mexico. Heredity 70:67.

47. Luo R, Liu B, Xie Y, Li Z, Huang W, Yuan J, He G, Chen Y, Pan Q, Liu Y, Tang J, Wu G, Zhang H, Shi Y, Liu Y, Yu C, Wang B, Lu Y, Han C, Cheung DW, Yiu SM, Peng S, Xiaoqian Z, Liu G, Liao X, Li Y, Yang H, Wang J, Lam TW,Wang J (2012) SOAPdenovo2: an empirically improved memory-efficient short-read de novo assembler. Gigascience 1(1):18.

48. Kim J, Larkin DM, Cai Q, Asan, Zhang Y, Ge RL, Auvil L, Capitanu B, Zhang G, Lewin HA,Ma J (2013) Reference-assisted chromosome assembly. Proc Natl Acad Sci U S A 110(5):1785–1790.

49. Künstner A, Hoffmann M, Fraser BA, Kottler VA, Sharma E, Weigel D,Dreyer C (2016) The Genome of the Trinidadian Guppy, Poecilia reticulata, and Variation in the Guanapo Population. Plos One 11(12):e0169087.

50. Kasahara M, Naruse K, Sasaki S, Nakatani Y, Qu W, Ahsan B, Yamada T, Nagayasu Y, Doi K, Kasai Y, Jindo T, Kobayashi D, Shimada A, Toyoda A, Kuroki Y, Fujiyama A, Sasaki T, Shimizu A, Asakawa S, Shimizu N, Hashimoto S, Yang J, Lee Y, Matsushima K, Sugano S, Sakaizumi M, Narita T, Ohishi K, Haga S, Ohta F, Nomoto H, Nogata K, Morishita T, Endo T, Shin IT, Takeda H, Morishita S,Kohara Y (2007) The medaka draft genome and insights into vertebrate genome evolution. Nature 447(7145):714–719.

51. Kitano J,Peichel CL (2012) Turnover of sex chromosomes and speciation in fishes. 94(3):549–558.

52. Meredith RW, Pires MN, Reznick DN,Springer MS (2011) Molecular phylogenetic relationships and the coevolution of placentotrophy and superfetation in Poecilia (Poeciliidae: Cyprinodontiformes). Mol Phylogenet Evol 59(1):148–157.

53. Mank JE, Promislow DE, Avise JC (2006) Evolution of alternative sex-determining mechanisms in teleost fishes. Biological Journal of the Linnean Society 87:83–93.

54. Pennell MW, Mank JE, Peichel CL (2018) Transitions in sex determination and sex chromosomes across vertebrate species. Mol Ecol 27(19):3950–3963.

55. Kondo M, Nanda I, Schmid M,Schartl M (2009) Sex Determination and Sex Chromosome Evolution: Insights from Medaka. Sexual Development 3(2-3):88-98.

56. Orzack SH, Sohn JJ, Kallman KD, Levin SA, Johnston R (1980) Maintenance of the three sex chromosome polymorphism in the platyfish, Xiphophorus maculatus. Evolution 34:663–672.

57. Kitano JU, Mori S, Peichel CL (2007) Phenotypic divergence and reproductive isolation between sympatric forms of Japanese threespine sticklebacks. Biological Journal of the Linnean Society 91:671–685.

58. Charlesworth D,Charlesworth B (2009) Sex differences in fitness and selection for centric fusions between sex-chromosomes and autosomes. Genetical Research 35(02):205.

59. Van Doorn GS, Kirkpatrick M (2010) Transitions between male and female heterogamety caused by sex-antagonistic selection. Genetics 186:629–645.

60. Reznick DN, Miles DB, Winslow S (1992) Life History of Poecilia picta (Poeciliidae) from the Island of Trinidad. Copeia 1992(3):782.

61. Russell ST, Ramnarine IW, Mahabir R,Magurran AE (2006) Genetic detection of sperm from forced copulations between sympatric populations of Poecilia reticulata and Poecilia picta. Biological Journal of the Linnean Society 88:397–402.

62. Ramsay C (2014) Mate Choice and Hybridization: Comparing the Endler’s Guppy (Poecilia wingei) and the Common Guppy (Poecilia reticulata).

63. Lohse M, Bolger AM, Nagel A, Fernie AR, Lunn JE, Stitt M,Usadel B (2012) RobiNA: A user-friendly, integrated software solution for RNA-Seq-based transcriptomics. Nucleic Acids Res. 40:622–627.

64. Kelley DR, Schatz MC, Salzberg SL (2010) Quake: quality-aware detection and correction of sequencing errors. Genome Biol 11:R116.

65. Chikhi R,Medvedev P (2014) Informed and automated k-mer size selection for genome assembly. Bioinformatics 30(1):31–37.

66. Kent WJ, Baertsch R, Hinrichs A, Miller W,Haussler D (2003) Evolution’s cauldron: duplication, deletion, and rearrangement in the mouse and human genomes. Proc Natl Acad Sci U S A 100(20):11484–11489.

67. Langmead B, Trapnell C, Pop M,Salzberg SL (2009) Ultrafast and memory-efficient alignment of short DNA sequences to the human genome. Genome Biol 10(3):R25.

68. Li H,Durbin R (2009) Fast and accurate short read alignment with Burrows-Wheeler transform. Bioinformatics 25(14):1754–1760.

69. Kim D, Langmead B,Salzberg SL (2015) HISAT: A fast spliced aligner with low memory requirements. Nat Methods 12(4):357–360.

70. Pertea M, Pertea GM, Antonescu CM, Chang TC, Mendell JT, Salzberg SL (2015) StringTie enables improved reconstruction of a transcriptome from RNA-seq reads. Nature Biotechnol 33:290–295.

71. Quinlan AR, Hall IM (2010) Genome analysis BEDTools: A flexible suite of utilities for comparing genomic features. Bioinformatics 26:841–842.

72. Flicek P, Amode MR, Barrell D, Beal K, Billis K, Brent S, …, Searle SMJ (2014) Ensembl 2014. Nucleic Acids Res. 42:749–755.

73. Anders S, Pyl PT, Huber W (2015) HTSeq--a Python framework to work with highthroughput sequencing data. Bioinformatics 31(2):166–169.

74. Robinson MD, McCarthy DJ, Smyth GK (2010) edgeR: a Bioconductor package for differential expression analysis of digital gene expression data. Bioinformatics 26(1):139–140.

75. Rahman A, Hallgrimsdottir I, Eisen M,Pachter L (2018) Association mapping from sequencing reads using k-mers. Elife 7.

76. Quinn A, Juneja P, Jiggins FM (2014) Estimates of allele-specific expression in Drosophila with a single genome sequence and RNA-seq data. Bioinformatics 30:2603–2610.

77. Zimmer F, Harrison PW, Dessimoz C,Mank JE (2016) Compensation of Dosage-Sensitive Genes on the Chicken Z Chromosome. Genome Biology and Evolution 8(4):1233–1242.

78. Koboldt DC, Zhang Q, Larson DE, Shen D, McLellan MD, Lin L, Miller CA, Mardis ER, Ding L,Wilson RK (2012) VarScanw 2: Somatic mutation and copy number alteration discovery in cancer by exome sequencing. Genome Res. 22:568–576.

79. Stevenson KR, Coolon JD, Wittkopp PJ (2013) Sources of bias in measures of allelespecific expression derived from RNA-seq data aligned to a single reference genome. BMC Genomics 14:536.

