## Supporting Information for "Extreme heterogeneity in sex chromosome differentiation and dosage compensation in livebearers"

### SI Results

**Table S1:** Sequencing results for each sample.

| Species (Treatment) | Sample no. (Sex) | Paired reads after trimming | % kept after trimming | Coverage |
|---|---|---|---|---|
| ***Poecilia wingei* (DNA-seq PE)** | 291 (F) | 222,019,309 | 97.7 | 77X |
| | 292 (F) | 209,095,391 | 92.6 | 72X |
| | 293 (F) | 244,778,587 | 92.3 | 85X |
| | 294 (M) | 221,308,140 | 92.9 | 76X |
| | 295 (M) | 245,199,642 | 93.3 | 85X |
| | 296 (M) | 214,802,737 | 93.2 | 74X |
| ***Poecilia picta* (DNA-seq PE)** | 247 (F) | 201,783,529 | 92.5 | 70X |
| | 248 (F) | 248,146,529 | 93.4 | 86X |
| | 265 (F) | 251,440,989 | 93.2 | 87X |
| | 266 (M) | 264,471,289 | 93.4 | 91X |
| | 267 (M) | 209,266,241 | 93.3 | 72X |
| | 268 (M) | 213,098,477 | 93.7 | 74X |
| ***Poecilia latipinna* (DNA-seq PE)** | 269 (F) | 242,950,245 | 93.6 | 83X |
| | 270 (F) | 186,547,462 | 92.7 | 64X |
| | 271 (F) | 194,577,608 | 92.7 | 67X |
| | 272 (M) | 235,795,174 | 93.3 | 81X |
| | 289 (M) | 229,757,997 | 93.4 | 79X |
| | 290 (M) | 232,391,653 | 93.0 | 80X |
| ***Gambusia holbrooki* (DNA-seq PE)** | 241 (F) | 217,994,173 | 93.8 | 75X |
| | 242 (F) | 193,263,881 | 93.5 | 67X |
| | 243 (F) | 229,309,343 | 93.3 | 79X |
| | 244 (M) | 195,792,613 | 93.4 | 68X |
| | 245 (M) | 194,586,542 | 93.6 | 67X |
| | 246 (M) | 220,591,540 | 93.4 | 76X |
| ***Poecilia wingei* (DNA-seq MP)** | 013 (F) | 80,809,424 | 58.0 | 23X |
| | 014 (F) | 76,562,926 | 58.1 | 22X |
| | 015 (F) | 77,120,163 | 58.5 | 22X |
| | 016 (M) | 75,360,153 | 56.4 | 22X |
| | 018 (M) | 80,705,804 | 57.9 | 23X |
| | 019 (M) | 83,808,049 | 58.8 | 24X |
| ***Poecilia picta* (DNA-seq MP)** | 013 (F) | 81,263,670 | 57.7 | 23X |
| | 014 (F) | 75,174,083 | 56.9 | 22X |
| | 015 (F) | 86,920,083 | 57.1 | 25X |
| | 016 (M) | 73,917,330 | 56.4 | 21X |
| | 018 (M) | 79,696,940 | 56.0 | 23X |
| | 019 (M) | 76,727,662 | 57.1 | 22X |
| ***Poecilia latipinna* (DNA-seq MP)** | 002 (F) | 87,479,612 | 56.1 | 25X |
| | 004 (F) | 87,085,262 | 56.8 | 25X |

| | | | | |
|---|---|---|---|---|
| | 005 (F) | 54,308,904 | 56.4 | 16X |
| | 006 (M) | 78,744,655 | 57.0 | 23X |
| | 007 (M) | 84,406,439 | 54.2 | 24X |
| | 012 (M) | 88,707,007 | 58.9 | 26X |
| | 002 (F) | 82,118,221 | 66.3 | 23.6 |
| | 004 (F) | 76,472,890 | 55.6 | 22.0 |
| ***Gambusia holbrooki*** **(DNA-seq MP)** | 005 (F) | 77,475,370 | 54.2 | 22.3 |
| | 006 (M) | 66,891,462 | 56.4 | 19.2 |
| | 007 (M) | 72,014,055 | 56.5 | 20.7 |
| | 012 (M) | 63,635,368 | 56.8 | 18.3 |
| | 201 (F) | 35,176,172 | 94.0 | - |
| | 202 (F) | 47,040,049 | 94.4 | - |
| ***Poecilia wingei*** **(RNA-seq)** | 203 (F) | 48,558,664 | 94.4 | - |
| | 265 (M) | 44,255,632 | 94.2 | - |
| | 266 (M) | 41,375,146 | 94.2 | - |
| | 267 (M) | 42,277,857 | 93.9 | - |
| | 282 (F) | 33,616,549 | 93.9 | - |
| | 284 (F) | 43,438,223 | 94.3 | - |
| ***Poecilia picta*** **(RNA-seq)** | 285 (M) | 45,953,612 | 94.3 | - |
| | 286 (M) | 39,836,450 | 94.0 | - |
| | 287 (M) | 43,314,678 | 94.1 | - |
| | 302 (F) | 48,435,135 | 94.0 | - |
| | 228 (F) | 48,056,489 | 94.3 | - |
| | 229 (F) | 34,836,324 | 94.3 | - |
| ***Poecilia latipinna*** **(RNA-seq)** | 230 (M) | 35,640,155 | 94.7 | - |
| | 231 (M) | 34,564,529 | 93.8 | - |
| | 232 (M) | 34,774,385 | 93.9 | - |
| | 288 (F) | 50,234,040 | 94.0 | - |
| | 204 (F) | 38,909,731 | 94.5 | - |
| | 205 (F) | 44,717,526 | 94.9 | - |
| ***Gambusia holbrooki*** **(RNA-seq)** | 206 (F) | 45,915,199 | 97.7 | - |
| | 207 (M) | 46,496,039 | 94.1 | - |
| | 208 (M) | 42,781,352 | 94.3 | - |
| | 281 (M) | 39,993,511 | 93.1 | - |

**Table S2:** Assembly statistics.

| Species | Total assembly length (Mb) | N50 (kb) | No. *de novo* scaffolds | No. RACA PCFs |
|---|---|---|---|---|
| ***P. wingei*** | 795.5 | 14.6 | 120,169 | 32 |
| ***P. picta*** | 782.2 | 150.6 | 9,640 | 102 |
| ***P. latipinna*** | 787.5 | 90.3 | 13,851 | 287 |
| ***G. holbrooki*** | 617.8 | 6.1 | 137,790 | 1,264 |

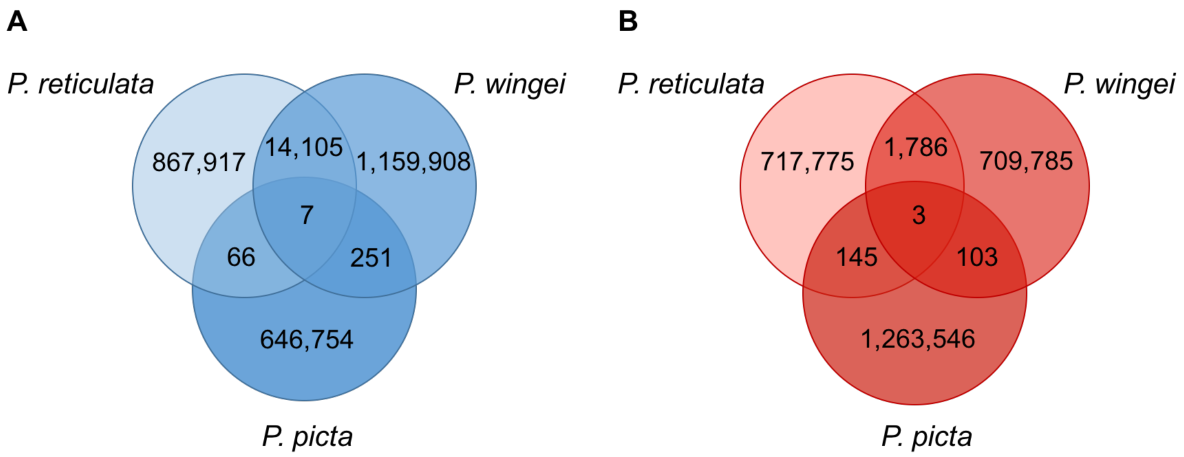


**Figure S1:** Number of shared *k*-mers across *P. reticulata*, *P. wingei* and *P. picta*. Species-specific and shared (A) male-unique *k*-mer (Y-mer) counts and (B) female-unique *k*-mer counts.

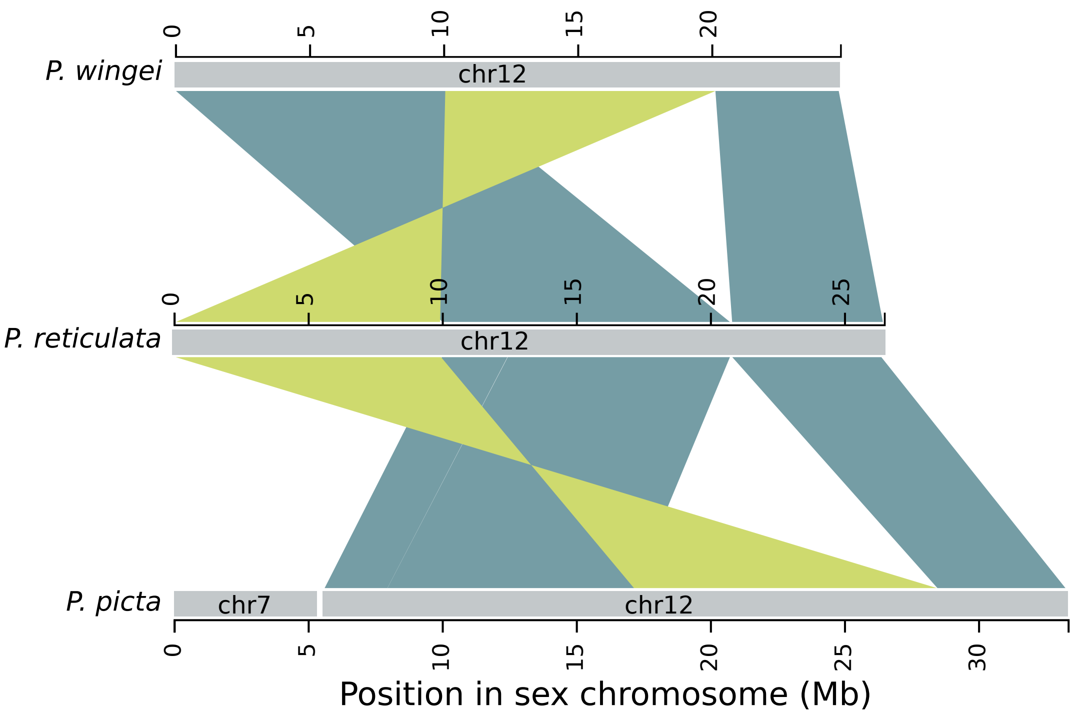


**Figure S2:** Synteny between the *P. reticulata*, *P. wingei* and *P. picta* sex chromosomes. Shown is the orientation of scaffolds mapping to *P. reticulata* chromosome 12. Regions showing the same orientation are shown in blue, inverted regions in green. The inversion is likely on the X chromosome rather than the Y as we obtain the same result when reconstructing chromosome fragments using read mapping data from female individuals alone.

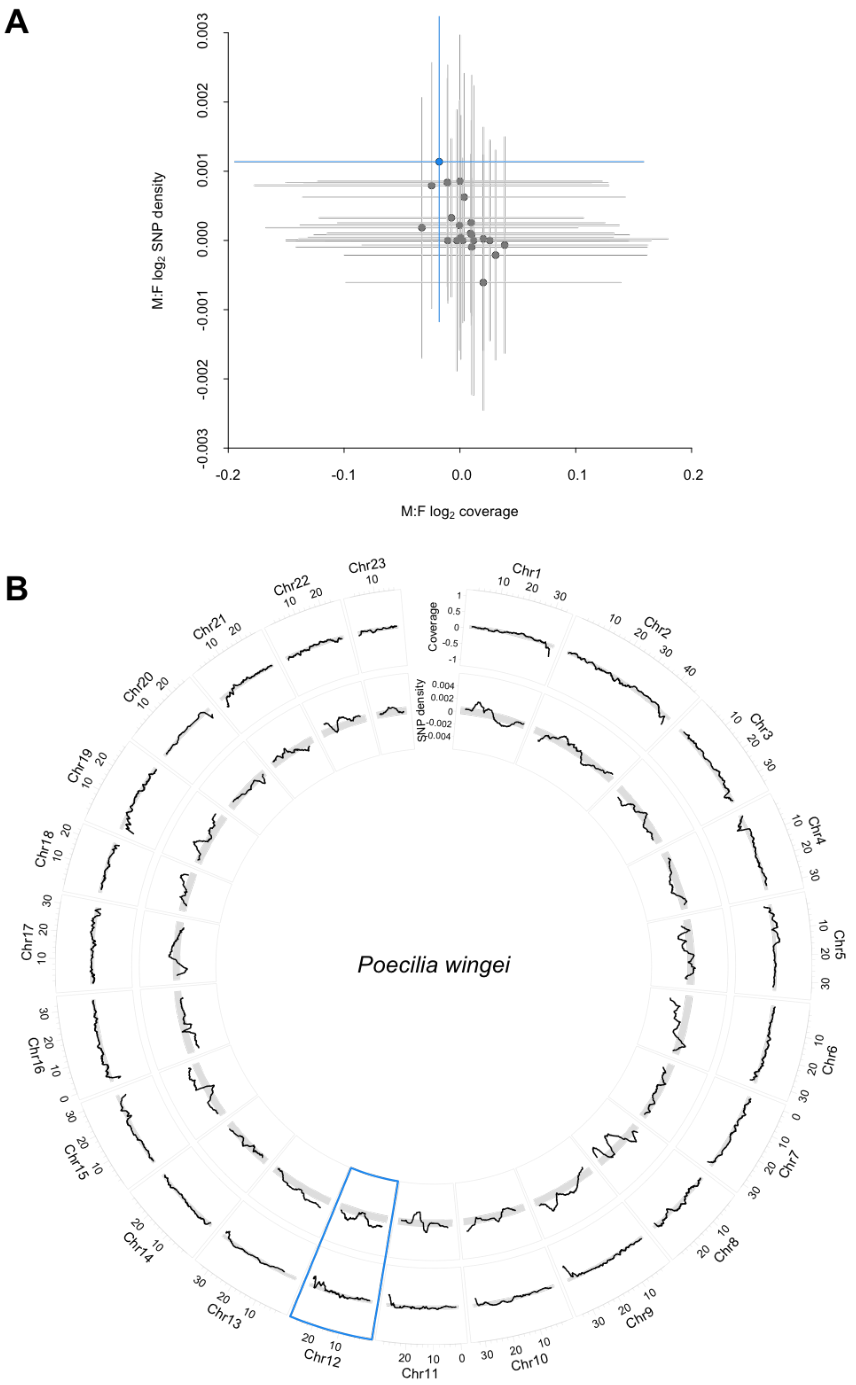


**Figure S3:** Coverage and SNP density differences between the sexes (M:F) across *P. wingei* chromosomes. (A) Average coverage and SNP density fold change for each chromosome. Shown in blue is chromosome 12, the sex chromosome in *P. wingei.* Interquartile ranges are represented by the vertical and horizontal lines. (B) Circos plot $log_2$ M:F coverage (outer ring) and M:F SNP density (inner ring) fold change moving average across each chromosome. Shown in grey are the 95% confidence intervals based on bootstrap estimates across the autosomes. Highlighted in blue is the XY sex chromosome in this species.

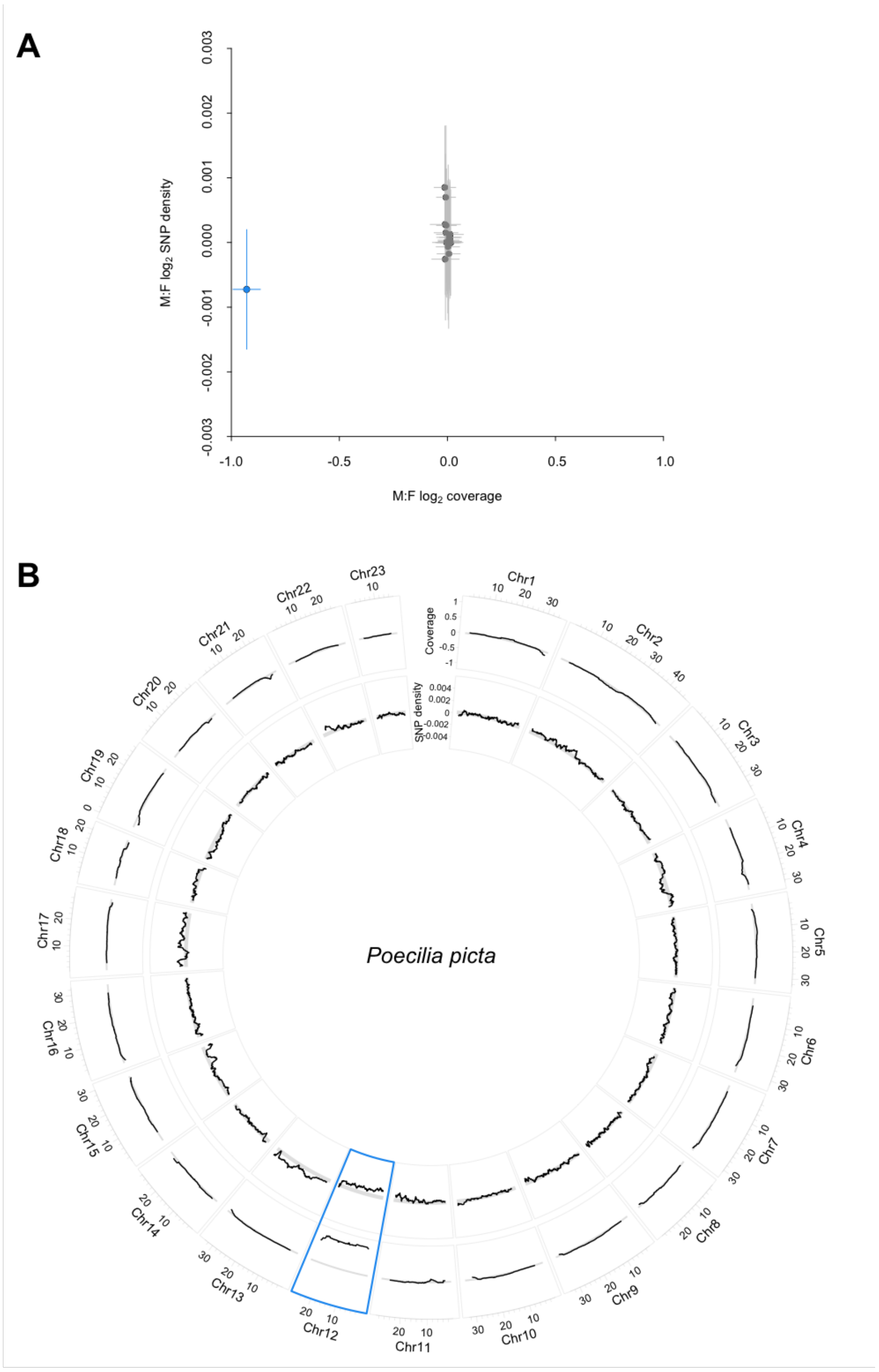


**Figure S4:** Coverage and SNP density differences between the sexes (M:F) across *P. picta* chromosomes. (A) Average coverage and SNP density fold change for each chromosome. Shown in blue is chromosome 12, the sex chromosome in *P. picta.* Interquartile ranges are represented by the vertical and horizontal lines. (B) Circos plot $\log_2$ M:F coverage (outer ring) and M:F SNP density (inner ring) fold change moving average across each chromosome. Shown in grey are the 95% confidence intervals based on bootstrap estimates across the autosomes. Highlighted in blue is the XY sex chromosome in this species.

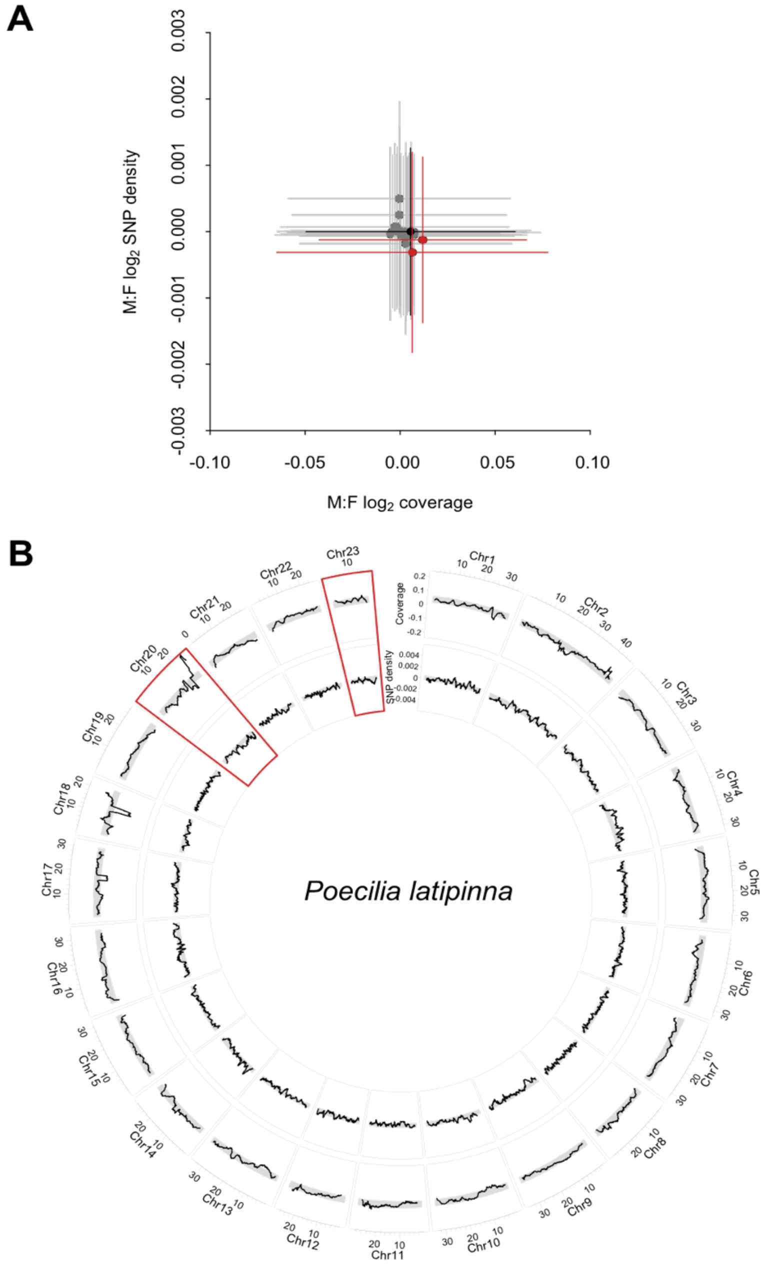


**Figure S5:** Coverage and SNP density differences between the sexes (M:F) across *P. latipinna* chromosomes. (A) Average coverage and SNP density fold change for each chromosome. Shown in red are the putative ZW sex chromosome candidates in this species, while chromosome 12, the sex chromosome in *P. reticulata, P. wingei* and *P. picta,* is shown in black. Interquartile ranges are represented by the vertical and horizontal lines. (B) Circos plot $log_2$ M:F coverage (outer ring) and M:F SNP density (inner ring) fold change moving average across each chromosome. Shown in grey are the 95% confidence intervals based on bootstrap estimates across the genome. Highlighted in red are the putative ZW sex chromosome candidates as identified in (A).

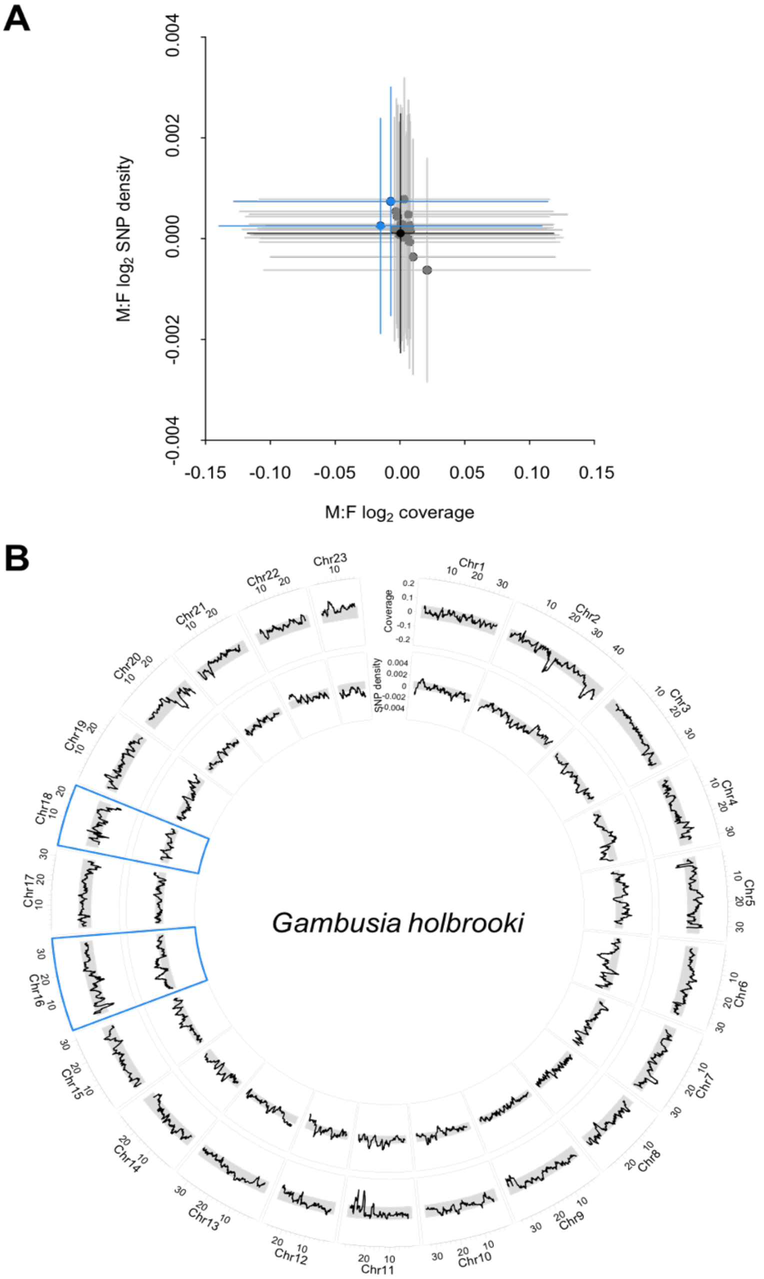


**Figure S6:** Coverage and SNP density differences between the sexes (M:F) across *G. holbrooki* chromosomes. (A) Average coverage and SNP density fold change for each chromosome. Shown in blue are the putative XY sex chromosome candidates in this species, while chromosome 12, the sex chromosome in *P. reticulata, P. wingei* and *P. picta,* is shown in black. Interquartile ranges are represented by the vertical and horizontal lines. (B) Circos plot of $log_2$ M:F coverage (outer ring) and M:F SNP density (inner ring) fold change moving average across each chromosome. Shown in grey are the 95% confidence intervals based on bootstrap estimates across the genome. Highlighted in blue are the putative XY sex chromosome candidates as identified in (A).

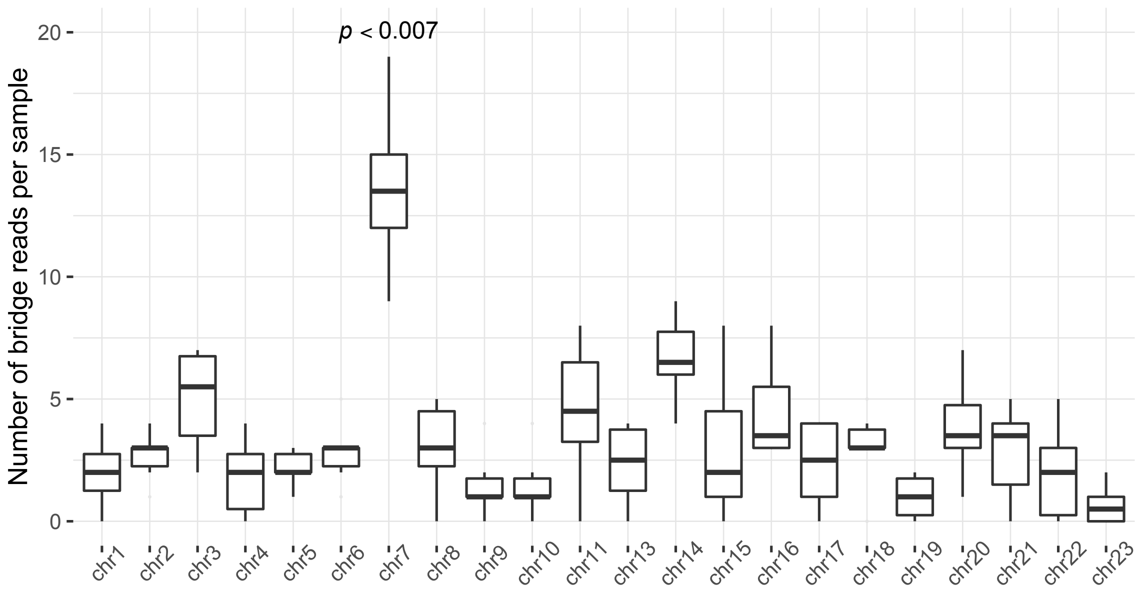


**Figure S7:** Number of bridge reads per individual supporting the fusion between the ancestral sex chromosome (chromosome 12) and each autosome in *P. picta*. Kruskal-Wallis, $p < 0.001$, significance based on pairwise Wilcoxon rank sum tests, all $p < 0.007$.
